# Quantitative Assays Reveal Cell Fusion at Minimal Levels of SARS-CoV-2 Spike Protein and Fusion-from-Without

**DOI:** 10.1101/2020.10.15.340604

**Authors:** Samuel A. Theuerkauf, Alexander Michels, Vanessa Riechert, Thorsten J. Maier, Egbert Flory, Klaus Cichutek, Christian J. Buchholz

## Abstract

Cell entry of the pandemic virus SARS-CoV-2 is mediated by its spike protein S. As main antigenic determinant, S protein is in focus of antibody-based prophylactic and therapeutic strategies. Besides particle-cell fusion, S mediates fusion between infected and uninfected cells resulting in syncytia formation. Here we present quantitative assay systems covering not only particle-cell and cell-cell fusion, but also demonstrating fusion-from-without (FFWO), the formation of syncytia induced by S-containing viral particles in absence of newly synthesized S protein. Based on complementation of split β-galactosidase and virus-like-particles (VLPs) displaying S protein, this assay can be performed at BSL-1. All three assays provided readouts with a high dynamic range and signal-to-noise ratios covering several orders of magnitude. The data obtained confirm the enhancing effect of trypsin and overexpression of angiotensin-converting enzyme 2 (ACE2) on membrane fusion. Neutralizing antibodies as well as sera from convalescent patients inhibited particle-cell fusion with high efficiency. Cell-cell fusion, in contrast, was only moderately inhibited despite requiring much lower levels of S protein, which were below the detection limit of flow cytometry and Western blot. The data indicate that syncytia formation as a pathological consequence in tissues of Covid-19 patients can proceed at low levels of S protein and may not be effectively prevented by antibodies.

## Introduction

The emergence of the so-called SARS-coronavirus-2 (SARS-CoV-2) is met with an unprecedented global scientific effort. As causative agent of the pandemic and the associated disease COVID-19, SARS-CoV-2 is a typical coronavirus, with a large spike protein (S) inserted in the membrane of the enveloped particle. Being the main surface-exposed antigen and the mediator of cell entry, S protein forms the most important target of current efforts to develop therapeutic drugs and prophylactic vaccines (Gioia *et al.*, 2020; Tang *et al.*, 2020).

As a typical class I viral fusion protein, S protein forms trimers with a globular head domain containing the receptor-binding site which contacts angiotensin-converting enzyme 2 (ACE2) as its entry receptor (Hoffmann *et al.*, 2020b; Lan *et al.*, 2020; Monteil *et al.*, 2020; Shang *et al.*, 2020b; Walls *et al.*, 2020; Yan *et al.*, 2020). Proteolytic cleavage separates the globular head (S1 domain) and the stalk domain (S2), which contains the hydrophobic fusion peptide at its N-terminus and the transmembrane domain towards the C-terminus. Priming of the fusion-competent state requires processing of two cleavage sites localized in close N-terminal proximity of the fusion peptide. These cleavage sites can be recognized by alternative proteases, thus explaining the high flexibility of the virus to adapt to various tissues. In particular, the S1/S2 site is cleaved by proprotein convertases like furin localized in the trans-golgi network during trafficking of the newly synthesized S protein to the cell surface. The S2’ site can be cleaved by the serine protease TMPRSS2 which is exposed on the surface of target cells and contacted when the virus binds to ACE2. In absence of TMPRSS2, virus particles are endocytosed and cleaved by cathepsins (Hasan *et al.*, 2020; Hoffmann *et al.*, 2020b; Hoffmann *et al.*, 2020a; Shang *et al.*, 2020a; Walls *et al.*, 2020). In cell culture, trypsin was described as further alternative protease able to activate membrane fusion (Ou *et al.*, 2020; Xia *et al.*, 2020; Zhang and Kutateladze, 2020). Thus, two alternative entry routes exist for SARS-CoV-2, notably determined by the availability of activating proteases (Hoffmann *et al.*, 2020b; Tang *et al.*, 2020). For both routes, membrane fusion is pH-independent.

Like other fusion proteins that are active pH-independently, S protein mediates not only fusion between the viral and the cellular membranes during particle entry, but also fusion of infected cells with uninfected cells. This process is mediated by newly synthesized S protein accumulating at the cell surface. The resulting syncytia are giant cells containing at least three, often many more nuclei. Cell-cell fusion is used by viruses such as human immunodeficiency virus (HIV) (Compton and Schwartz, 2017; Bracq *et al.*, 2018), measles virus (MV) (Griffin, 2020), or herpesvirus (Cole and Grose, 2003) to spread in a particle-independent way. The resulting syncytia are documented as pathological consequence detectable in various tissues such as lung (MV), skin (herpesvirus) or lymphoid tissues (HIV). In brain, cell-to-cell transmission via hyperfusogenic F proteins constitutes a hallmark of MV-caused encephalitis as a fatal consequence of acute MV infections manifesting years later (Ferren *et al.*, 2019).

Besides particle-cell and cell-cell fusion, a third membrane fusion process mediated by viral spike proteins has been designated fusion-from-without (FFWO) (Roller *et al.*, 2008). FFWO results in syncytia in absence of newly expressed fusion protein. In presence of a sufficient concentration of particle-associated fusion protein, adjacent cells are fused e.g. by bound HIV or herpesvirus particles either directly or after uptake of spike protein and its presentation at the cell surface (Clavel and Charneau, 1994; Melikyan, 2014).

Here, we investigated the competence of SARS-CoV-2 S protein for these three membrane fusion processes. For each of them, we established quantitative assays relying on expressed S protein, thereby avoiding work with infectious virus at BSL-3 safety level. The data reveal a strong membrane fusion activity of the S protein, demonstrate syncytia formation even at undetectable levels of S protein, and fusion-from-without. Examination of sera from convalescent Covid-19 patients revealed potent neutralizing capacity against particle-cell fusion, but only moderate or low activity against cell-cell fusion.

## Methods

### Expression plasmids

An expression plasmid for codon-optimized SARS-CoV-2 S with a C-terminal HA tag (pCG-SARS-CoV-2-S-HA) was kindly provided by Karl-Klaus Conzelmann (Hennrich *et al.*, 2020).The Plasmid pCG-SARS-CoV-2-SΔ19 encoding the SΔ19 variant was generated by PCR amplification of the truncated S sequence, inserting PacI and SpeI restriction sites as well as a stop codon by PCR with primers 5’-TTATTAATTAAATGTTCGTGTTTCTGGTG-3’ and 5’-TATACTAGTTCTAGCAGCAGCTGCC-3’. The PCR fragment was inserted into the pCG backbone by restriction cloning. The lentiviral transfer vector plasmid pCMV-LacZ was generated by amplifying the lacZ coding sequence under CMV promotor control, simultaneously inserting a SbfI restriction site for subcloning into pSEW via EcoRI/Bsu36I (Funke *et al.*, 2008). Expression plasmids pCMV-α and pCMV-ω encoding the α and ω parts of β-galactosidase have been described previously (Holland et al. 2004). The expression plasmid for N-terminally myc-tagged hACE2 (pCMV3-SP-Myc-ACE2) was purchased from Sino Biological (HG10108-NM).

### Cell culture and transfection

HEK-293T (Lenti-X 293T, Takara Bio) and Vero E6 (ATCC) cells were cultured in Dulbecco’s Minimal Essential Medium High Glucose (Sigma, D6546) supplemented with 10% FBS (Sigma, F7524, lot BCCB7222) and 1x L-glutamine (Sigma, G7513) (i.e. DMEM complete). HEK-293T cells were subcultured twice a week at ratios between 1:8 and 1:10 using 0.25% trypsin in 1 mM EDTA - PBS without Ca^2+^ or Mg^2+^. MRC-5 cells (ATCC) were cultured in Minimal Essential Medium Eagle (Sigma, M2414) supplemented with 10% FBS, 1x L-glutamine and 1x non-essential amino acids (Gibco, 11140-035). Calu-3 cells (AddexBio, C0016001, lot 0179286) were cultured in Minimal Essential Medium Eagle (Sigma, M2414) supplemented with 10% FBS, 1x L-glutamine, 1x non-essential amino acids (Gibco, 11140-035) and 1x sodium pyruvate (Gibco, 11360-070). MRC-5 and Calu-3 were subcultured with trypsin in EDTA-PBS every two weeks at ratios between 1:2 and 1:3, medium was exchanged twice weekly.

For transfection of HEK-293T in T75 flasks, two mixes were prepared, each in 1 mL plain DMEM High Glucose, one containing 60 μL of 18 mM TA-Trans (polyethyleneimine) and the other 15 μg plasmid DNA. When the amount of S protein encoding plasmid was varied, pCDNA3.1(+) was added to compensate for the overall plasmid quantity. The mixtures were combined and incubated together for at least 20 min. Medium on cells was replaced by 4.3 mL of 1.5 x growth medium (15% FBS, 1.5 x L-Glutamine) and 2 mL transfection mix were added. Cells were incubated for 4-8 h before medium was replaced by 10 mL of fresh growth medium.

### Production of LVs and VLPs

For the production of LV particles and VLPs, we adapted our established protocol for the generation of LVs pseudotyped with paramyxoviral glycoproteins (Bender *et al.*, 2016). In brief, HEK-293T cells were transfected with the envelope plasmid pCG-SARS-CoV-2-SΔ19, the transfer vector plasmid pCMV-LacZ and the packaging plasmid pCMVd8.9 in a 35:100:65 ratio. For VLPs, the transfer vector plasmid was omitted, but the total amount of plasmid and the ratio of envelope to packaging plasmid retained. Supernatants harvested 48 hours after transfection were clarified by 0.45 μm filtration. Particles were purified and concentrated 333-fold into PBS from the clarified supernatant by overnight centrifugation over a 20% sucrose cushion at 4500 x g and 4°C. Concentrated stocks were frozen to −80°C before use, and aliquots used once after thawing. Particle numbers were determined by nanoparticle tracking analysis using the NanoSight300 system (Malvern).

### Transduction and neutralization

2×10^4^ cells/ well were seeded into flat-bottom 96-well plates in complete growth medium. On the next day, 0.2 μL/well of SΔ19-LV or VSV-LV stock was added in complete growth medium. For neutralization, antibody and sera were incubated with the vector stock at concentrations of 40 μg/ mL and 50% v/v, respectively, in a final volume of 50 μL/well for 30-60 min at 37°C. Medium was aspirated from the seeded cells and replaced by 50 μL/well of the transduction mix. The next morning, medium was exchanged for 100 μL/well of fresh complete growth medium. Transduction efficiencies were determined three days after vector addition by luminescence readout of cellular galactosidase activity.

### Sera & S-Neutralizing Antibodies

Sera donation with informed consent was approved by an ethics vote from the local committee at Frankfurt University Hospital. Sera were from two convalescent patients who had been diagnosed with SARS-CoV-2 by PCR from throat swabs approximately 4 months prior to donation. Both had experienced only mild symptoms. A commercially available pool of human off-the-clot sera (PAN Biotech, P30-2701) served as a negative control. A commercially available neutralizing antibody against SARS-CoV-2 (Sino Biological, 40592-R001) and the corresponding normal control (Sino Biological, CR1) served as a positive control.

### Fusion assays and neutralization

HEK-293T cells were transfected as described above, in the T75 format. Two days after transfection, cells were detached by incubation in 0.25% trypsin - 1 mM EDTA - PBS for 10-15 min. Cells were characterized with regards to count and viability using the Luna-Fl cell counter and acridine orange/propidium iodide dye (Logos Bio). Cells were pelleted at 300 x g for 5 min and resuspended in complete growth medium to yield a cell density of 5×10^4^ cells/20 μL. Cocultures were set up in V-bottom plates at 10^5^ cells/well. Cells were pelleted (300 x g, 30 s at start) just before transfer to the incubator. For FFWO, VLPs were diluted to desired concentrations in complete growth medium and added to cocultures at 20 μL/well by thorough pipetting prior to the 30 s centrifugation. For neutralization of FFWO, 5×10^8^ particles/well were incubated with antibodies or sera for 30 min in the incubator before addition to the coculture. For the neutralization of particle-free cell fusion, effector cells were pre-incubated with antibodies or sera in a volume of 40 μL/well (i.e. 5×10^4^ cells/well) for 30 min in the incubator, retaining the inhibitor concentrations specified above. After pre-incubation, effector cells were mixed with target cells in a total culture volume of 60 μL/well.

### Luminescense readout

Activity of the β-galactosidase reporter was quantified using the Galactostar assay kit (Thermo, T1012). At the assay endpoint (3 days post transduction or 20 h after coculture setup), cultures were lysed: Cocultures in V-bottom plate were pelleted at 300 x g, 5 min. Supernatant was removed completely, 50 μL/well of lysis buffer was added and plates were agitated on an orbital shaker at 450 rpm at room temperature for 10 min. Plates were then frozen to −80°C. For the luminescence readout, samples were equilibrated to room temperature and mixed by orbital shaking at 750 rpm for 2 min. 10 μL/well of lysate was added to 50 μL/well of substrate working dilution

(prepared according to manufacturer’s instructions) in a white plastic flat-bottom plate and mixed by orbital shaking at 750 rpm for 2 min. After 30-60 min incubation at room temperature in the dark, luminescence was measured on an Orion II plate luminometer (Berthold Systems) with an exposure time of 0.1 s/well.

### Immunofluorescence staining and laser scanning microscopy

Vero E6 cells constitutively expressing GFP were generated by LV mediated transduction and subsequent puromycin selection. HEK-293T cells were cotransfected with RFP and the SΔ19 or full-length S protein. 5×10^4^ transfected HEK-293T cells were seeded in chamber slides (Thermo, 177402) and left to attach overnight. On the next day 5×10^4^ Vero-GFP cells were added and cocultured for 7 h. Cells were fixed in 4% PFA, permeabilized with 0.5% Triton X-100 in PBS and blocked with 1% BSA/PBS for 15 min. Subsequently, cells were stained with Phalloidin-Atto633 (1:500, Sigma 68825) and HOECHST3342 (1:10,000, Sigma B2261) for 1h at RT before being imaged on an SP8 Lightning laser scanning microscope (Leica) with a HC PL APO CS2 40x/1.30 lens.

### Western Blot

Cells were detached by trypsin treatment, counted with a hemocytometer and lyzed in RIPA buffer (50 mM Tris/HCL pH 8.0, 150 mM NaCl, 1% NP-40, 0.5% sodium deoxycholate and 0.1% SDS) supplemented with a protease inhibitor cocktail (Roche, 05892970001). Cell lysates were incubated for 10 min at 95°C in 4x sample buffer (240 mM Tris/HCL pH 6.8, 8% SDS, 40% glycerin, 0.2% bromphenol blue, 20% β-mercaptoethanol). Vector particles were denatured for 10 min at 95°C in 2x Urea buffer (200 mM TRIS/HCl pH 8.0, 5% SDS, 8 M Urea, 0.1 mM EDTA, 2.5% DTT, 0.03% bromphenol blue) (Münch *et al.*, 2011). Samples were electrophoretically separated on a 10% polyacrylamide gel and blotted onto nitrocellulose membranes (Amersham, 10600004). The lower part of the membranes were incubated with mouse anti-p24 (Clone 38/8.7.47, 1:1000, Gentaur) and the upper part with mouse anti-SARS-CoV-2 spike (Clone: 1A9, 1:1000, GeneTex) overnight at 4°C. Subsequently, the membranes were incubated with the secondary antibody rabbit anti-mouse conjugated to horseradish peroxidase (Dako, 1:2000) for 90 min at RT. Luminescence signals were detected on the chemiluminescence reader MicroChemi (DNR) after adding ECL Western Blotting Substrate (Thermo, 32106).

### Flow cytometry

10^5^ HEK-293T effector cells used for fusion assays were stained for surface expression of S protein. Cell suspensions were washed twice in wash buffer (2% FCS, 0.1% sodium azide, 1 mM EDTA in PBS). S protein was specifically stained with the mouse IgG1 anti-SARS-CoV-2 Spike (Clone: 1A9, GeneTex, 1 μL/10^5^ cells in 100 μL) antibody for 45 min at 4°C followed by the incubation with the secondary antibody anti-IgG1-PE (REA1017, Miltenyi Biotec, 1 μL/10^5^ cells in 100 μL) for 30 min at 4°C. Viability of the cells was assessed using the fixable viability dye eFluor780 (eBioscience, 1:1000). Finally, cells were fixed in 1% PFA and analyzed by flow cytometry using the MACSQuant Analyzer 10x (Miltenyi Biotec).

### Statistical Analysis

All statistical analyses were carried out in GraphPad Prism version 8.4.2. Luminescense data and flow cytometry MFIs were assumed to be lognormally distributed. Accordingly, tests were performed on log-transformed data which were assumed to be normally distributed. Data generated from the same batch of transfected cells (i.e. from the same biological replicate) were handled as matched data. For all repeated measures tests, sphericity was assumed. For particle and inhibitor titrations, five-parameter asymmetric sigmoidal curves were fitted to the data. If not otherwise indicated, symbols represent means of technical triplicates, bars represent means (geometric means for non-log-transformed data) of biological replicates and error bars represent 95 % CIs. Differences of population means were quantified by repeated measures 2-way ANOVA (one-way ANOVA, where indicated) on log-transformed data and Tukey’s multiple comparisons test. Select multiplicity-adjusted p-values are reported.

## Results

### Spike protein mediated particle entry

To follow particle entry mediated by S protein, we set out to generate potent lentiviral vectors (LVs) pseudotyped with S protein to set up an assay with a high signal-to-noise ratio. Codon-optimized S protein, full length or carrying the previously described C-terminal truncation (SΔ19), were used for pseudotyping (Ou *et al.*, 2020). LVs transferring the β-galactosidase gene *lacZ* were generated by transient transfection of HEK-293T cells and subsequently purified and concentrated (Fig. 1A). Western blot analysis of LV batches revealed a stronger spike signal relative to p24 for SΔ19, demonstrating its better incorporation into LV particles (Fig. 1B). SΔ19-LVs were titrated on cell lines frequently used in coronavirus research, i.e. Vero E6, MRC-5, Calu-3, HEK-293T and HEK-293T overexpressing ACE2 (293T-ACE2). On all cell lines included, the luminescense signal increased linearly with the amount of vector added (Fig. 1C). In contrast to LVs pseudotyped with the G protein of vesicular stomatitis virus (VSV-LV), SΔ19-LV did not reach a plateau in the signal, indicating that only a subsaturating fraction of the cells was transduced. However, transduction rates with SΔ19-LV increased more than 100-fold upon overexpression of hACE2 on HEK-293T cells reaching a signal to noise ratio of more than 2000. VSVG-LV mediated gene delivery was not affected by overexpression of hACE2 (Fig. 1D). Remarkably, with a saturating dose of SΔ19-LV, a similar luminescense signal was reached on 293T-ACE2 cells as with VSV-LV (Fig. 1C).

**Figure 1:**
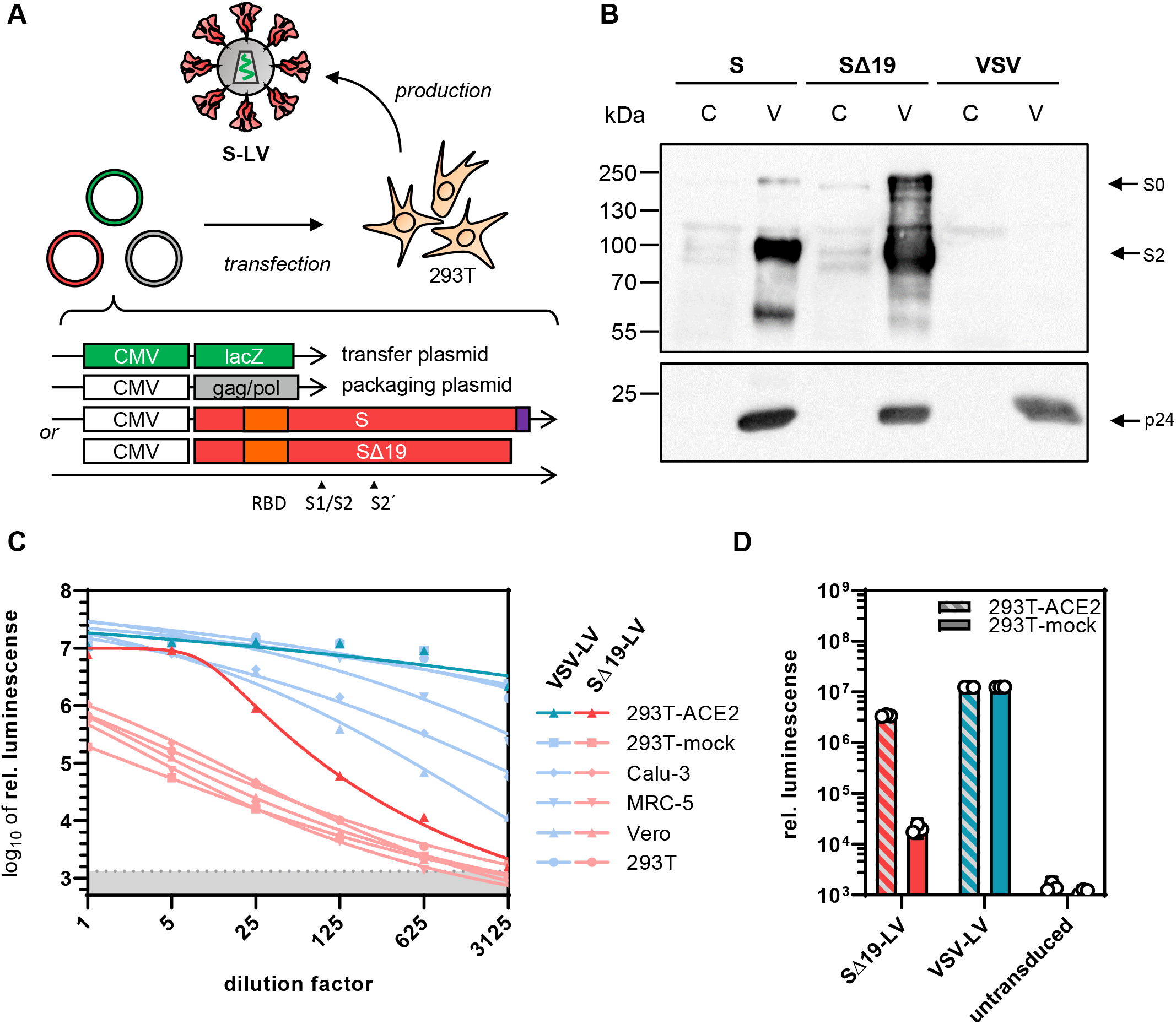
Spike mediated particle entry. (**A**) Generation of pseudotyped lentiviral vectors. Second generation LVs pseudotyped with S protein were generated by transfection of HEK-293T cells with a packaging plasmid encoding HIV-1 gag/pol, a transfer vector plasmid with a lacZ reporter gene and one of two envelope plasmids encoding codon-optimized SARS-CoV-2 S with or without (SΔ19) the 19 C-terminal amino acids. The C-terminal endoplasmic reticulum retention signal (purple) and the receptor binding domain (RBD, orange) are indicated. (**B**) Incorporation of S protein into LVs determined by Western blotting. S-LV and SΔ19-LV particles (V) and lysates of their producer cells (C) were stained for the presence of S protein (top) and p24 as particle loading control. Top blot was exposed for 30 s, bottom blot for 5 s. (**C**) Gene transfer activities on the indicated cell lines. The indicated dilutions of 5 μl vector stock of SΔ19-LV or VSV-LV were added to the cells. Cell lysates were prepared three days after vector addition and lacZ reporter activity was quantified as a luminescence readout. Symbols represent means of technical triplicates. Grey shaded area indicates 95% CI of signals from untransduced cells (blanks). (**D**) Effect of ACE2-overexpression on reporter transfer. 293T cells transfected with ACE2 expression plasmid or mock plasmid were incubated with 0.2 μL of SΔ19-LV or VSV-LV. Cell lysates were prepared three days after vector addition and reporter activity was quantified as a luminescence readout. Bars represent geometric means of technical triplicates ±95% CIs.

### Quantifying cell fusion mediated by SARS-CoV-2 S protein

Upon transfection, SARS-CoV-2 S protein showed a remarkable fusogenic activity. When transfected HEK-293T cells producing SΔ19-LV were detached and cocultured in a 1:1 ratio with Vero E6 cells, plates were covered by large syncytia, each containing at least 10, and, up to 100 nuclei, formed overnight (Suppl. Fig. 1A). To examine this further, the full-length and truncated forms of S were overexpressed in HEK-293T and cocultures with Vero E6 target cells were imaged by confocal laser scanning microscopy (Suppl. Fig. 1B). Both, S and SΔ19 protein, induced many large syncytia characterized by cytoskeletal rearrangement, clustering of more than five nuclei and colocalization of the RFP and GFP reporter dye signals (Suppl. Fig. 1C).

To quantify the cell-cell fusion mediated by S protein, an α-complementation assay based on β-galactosidase previously used to evaluate HIV-mediated fusion (Holland *et al.*, 2004) was adapted to the S protein. Upon cell-cell fusion, complementation inside the syncytium results in active enzyme that can be quantified by converting substrate in a luminescence reaction (Fig. 2A). In a first step, the minimal amount of S protein required to result in a detectable signal was determined. Effector cells were transfected with varying quantities of S protein encoding plasmid ranging and S protein surface expression was followed by flow cytometry. S protein expression was still clearly detectable with 75 ng of plasmid, while the mean fluorescence intensity (MFI) at 7.5 ng plasmid was indistinguishable from background (Fig. 2B). Notably, this low level of S protein was still sufficient to induce significant cell-cell fusion, when the assay was allowed to proceed over-night, highlighting the potent fusogenic activity of S protein (Fig. 2C). With higher amounts of plasmid, three hours incubation were sufficient to obtain signals more than 10-fold above background. Interestingly, when cells were detached prior to coculture with EDTA only instead of trypsin-EDTA, the cell fusion activity decreased by at least one order of magnitude, and only overnight incubation yielded signals significantly over background (Fig. 2D). When target cells overexpressed ACE2 and were detached with trypsin, the maximum extend of the fusion signal increased by another order of magnitude (Fig. 2D). Now, the signal-to-noise ratio reached two orders of magnitude already after one hour incubation including the samples expressing the lowest amounts of S protein. This highlights the dependence of S protein activity on the presence of ACE2. Under optimal conditions and with ACE2 overexpressing cells, a signal-to-noise ratio of 2.8 orders of magnitude was reached.

**Figure 2:**
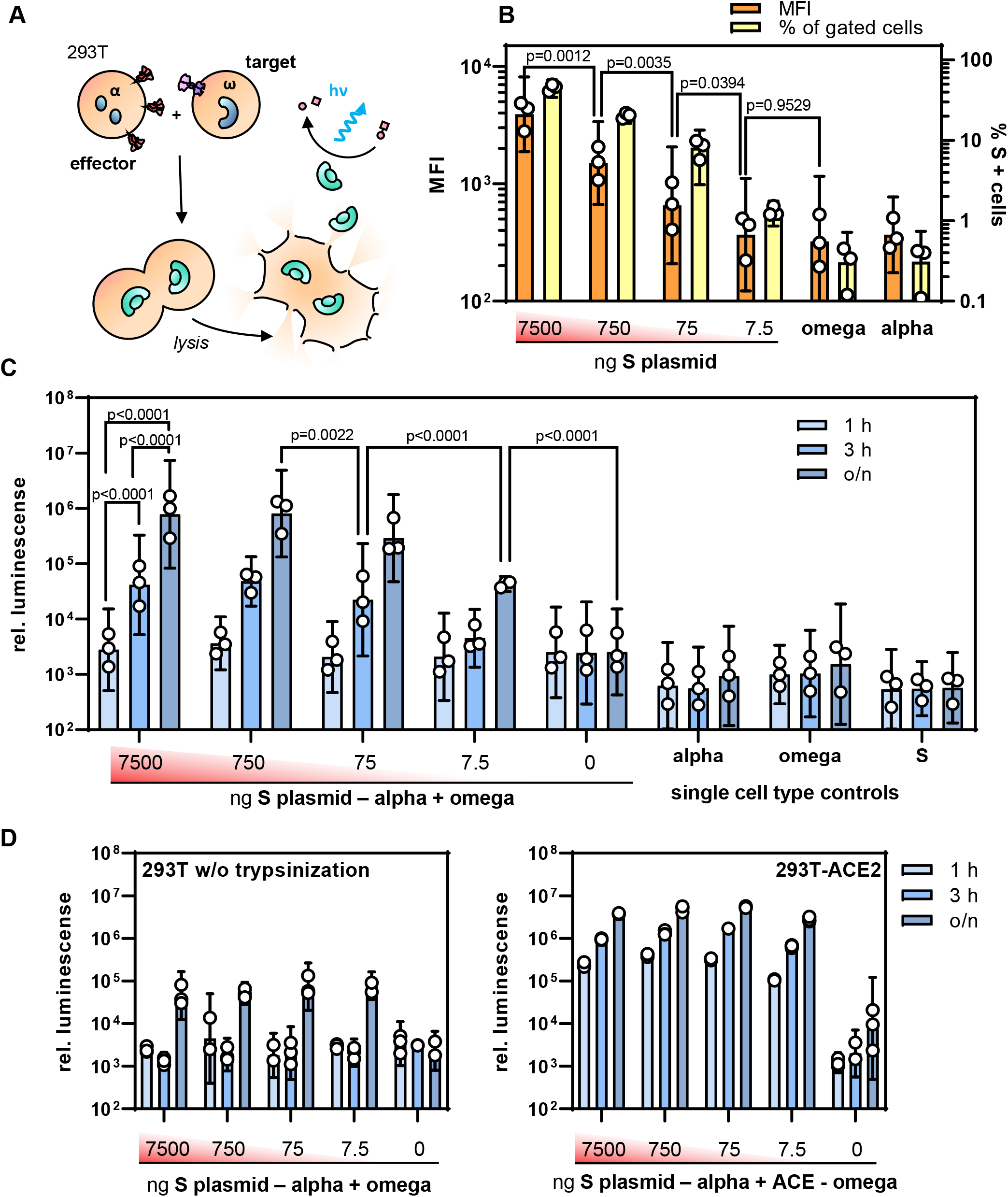
Quantifying S protein mediated cell-cell fusion. (**A**) Principle of quantifying cell-cell fusion. Effector cells expressing S protein and the α-fragment contain active β-galactosidase when they fuse with target cells expressing ACE2 and the ω-fragment. (**B-C**) Effector cells transfected with different amounts of S-protein expression plasmid (7500-7.5 ng/T75 flask) were assessed for S protein expression by flow cytometry (**B**) and then applied in the fusion assay (**C**). (**B**) Mean fluorescence intensity (MFI, orange bars) and the percentage of S-positive cells (yellow bars) were determined in flow cytometry. Bars represent means ± 95% CIs, n=3. P-values are from one-way ANOVA. (**C**) Activities of β-galactosidase in cocultures of effector and target cells in absence of ACE2 overexpression. Effector cells were transfected with the indicated amounts of S protein encoding plasmid, detached with trypsin and cultivated for the indicated time periods with target cells which were transfected with the ω-fragment encoding plasmid only. Bars represent means ± 95% CIs, n=3. (**D**) Influence of proteolytic processing and ACE2 overexpression on cell fusion activity. Left panel: Effector cells were detached without trypsin using 5 mM EDTA in PBS. Right panel: Target cells were co-transfected with the ω-fragment and ACE2 encoding plasmids. Bars represent means ± 95% CIs, n=3.

### Fusion-from-without mediated by S protein

To examine whether S protein present on the surface of viral particles can mediate FFWO, we generated LV-based virus-like-particles (VLPs) having the SΔ19 protein incorporated. Western blot analysis confirmed that similar levels of SΔ19 protein were present in VLPs and SΔ19-LV particles (Fig. 4E). To quantify FFWO, cocultures of 293T-ACE2 cells transfected with the plasmids encoding the alpha or the omega fragment of β-galactosidase were incubated with different amounts of VLPs to induce cell fusion and thus complementation (Fig. 3A). At a dose of 5000 VLPs per cell, a substantial fusion signal more than one order of magnitude over background was obtained (Fig. 3B). This signal increased further with increasing particle numbers, reaching a plateau at 5×10^4^ particles per cell, with a signal-to-noise ratio of 2.7 orders of magnitude (Fig. 3B). This FFWO activity of the S protein was again strongly dependent on ACE2 overexpression since no significant fusion activity was observed in absence of ACE2 (Suppl. Fig. 2). When the VLPs were treated with 2 mg/mL trypsin, the fusion signal increased by approximately 5.7-fold after 30 min of incubation. The chosen incubation period was critical for this increase, since prolonged incubation with trypsin was not beneficial (Fig. 3C). This correlated well with processing of the S0 protein into S2 for short trypsin exposure while prolonged digestion resulted in loss of S2 (Fig. 3D).

**Figure 3:**
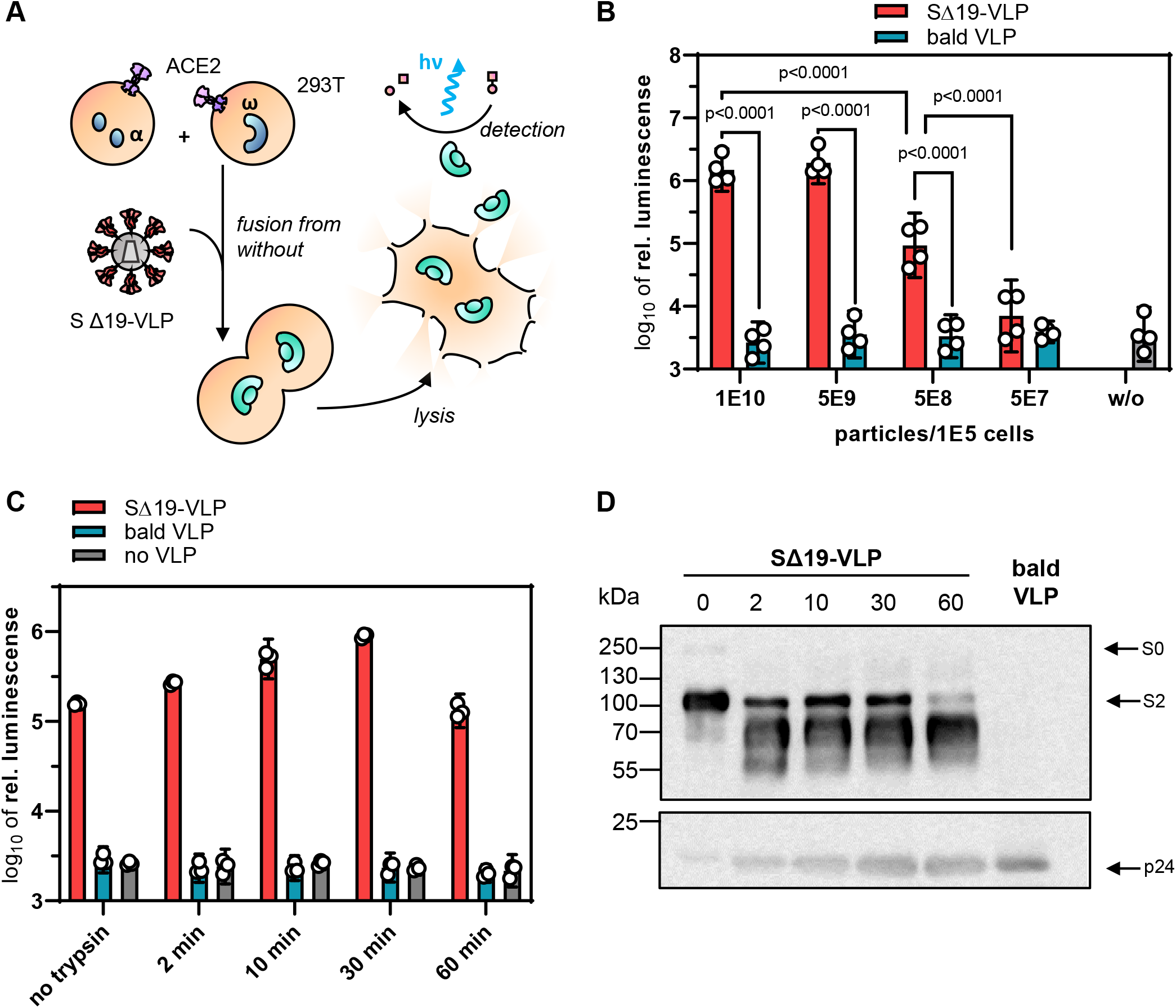
S protein mediated fusion-from-without. (**A**) Principle of the assay. HIV-1 derived SΔ19-VLPs were added as effector triggering fusion from without (FFWO) of cocultures of ACE2 overexpressing HEK-293T cells transfected with plasmids encoding the α-fragment or the ω-fragment of β-galactosidase. Cocultures were lysed and galactosidase activity of the lysates determined in luminescence reactions. (**B**) The indicated numbers of SΔ19-VLPs or bald VLP, produced without the S protein, were added to the coculture. Activities were quantified after overnight incubation. Bars represent means ± 95% CIs, n=4. (**C-D**) Effect of proteolytic processing on the induction of FFWO. 5×10^8^ SΔ19-VLPs, bald VLPs or no VLPs were incubated in 2 mg/mL trypsin for the indicated time periods before addition to cocultures (10^5^ cells). Luminescence was quantified after overnight cultivation. (**C**). Bars represent means of technical triplicates ± 95% CIs. (**D**) Processing of the S protein after trypsin treatment of the VLPs for indicated times by Western blotting for S and p24 proteins.

**Figure 4:**
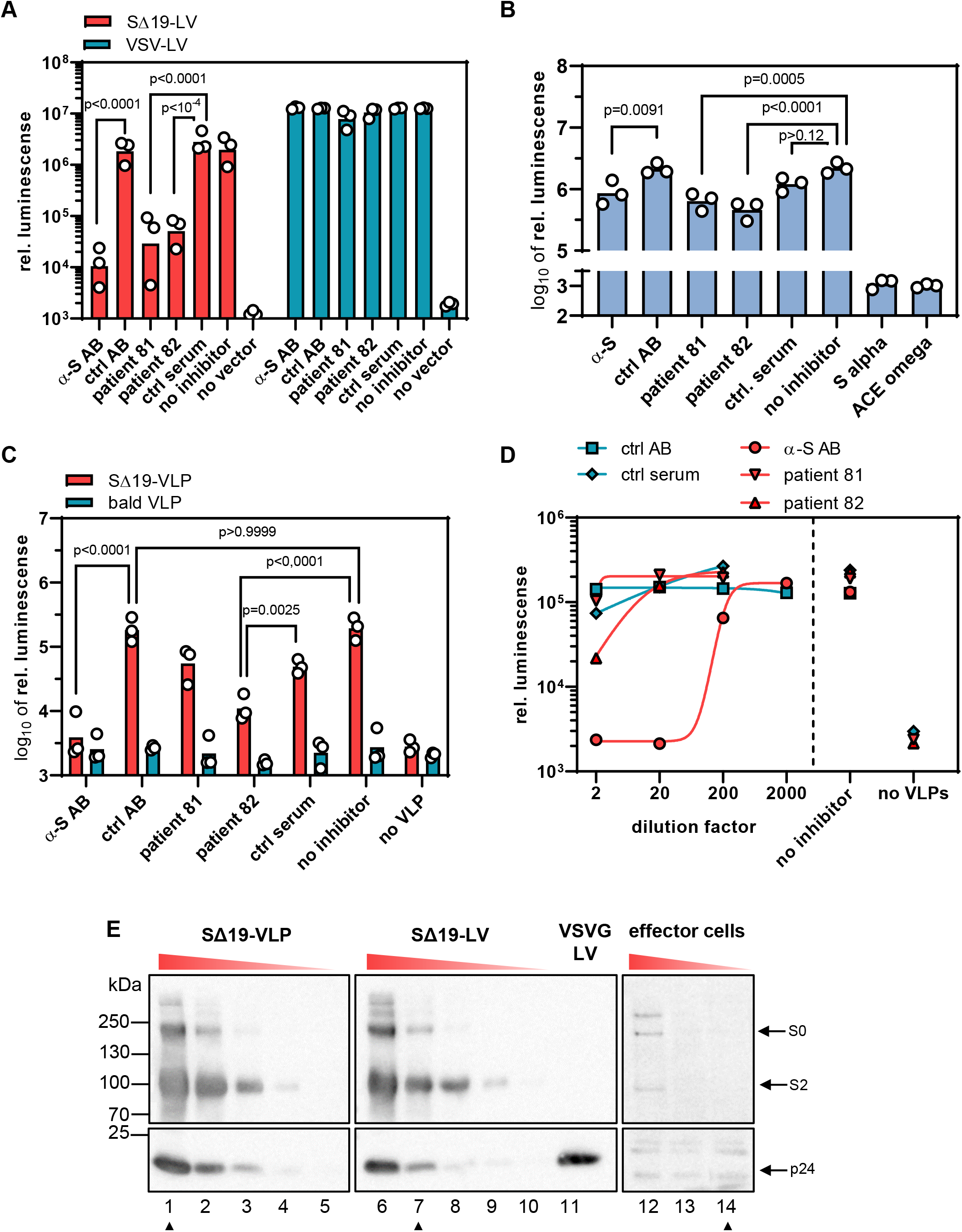
Antibody mediated neutralization of membrane fusion. The neutralizing activities of an S-protein specific antibody and sera from two convalescent Covid-19 patients were determined against S-protein mediated particle entry (**A**), cell-cell fusion (**B**) and FFWO (**C-D**). Bars represent means, n=3. P-values are from two-way (**A, C**) or one-way (**B**) ANOVA. (**A**) 0.2 μL of SΔ19-LV or VSV-LV were incubated for 30 min with the indicated antibodies or sera before being added to HEK-293T cells transfected with ACE2 expression plasmid. Cell lysates were prepared 3 days after vector addition and reporter activity was quantified as luminescence readout. (**B**) Effector cells coexpressing S protein (7.5 ng S plasmid per T75) and the α-fragment were incubated for 30 min with antibodies or sera before being added to target cells cotransfected with ω-fragment and ACE2 expression plasmids Bars represent means, n=3. (**C**) 5×10^8^ SΔ19-VLPs or bald VLPs were incubated for 30 min with antibodies or sera before being added to the cocultures of α/ACE2 and ω/ACE2 cells. Luminescence was quantified after overnight incubation. Select multiplicity-adjusted p-values are reported. (**D**) Neutralizing capacity of the indicated dilutions of antibodies and sera on FFWO. Dilution factors apply to pure serum 80 μg/mL for the antibodies. (**E**) Semi-quantitative Western blots comparing S protein dose applied to the three fusion assays as SΔ19-VLPs, SΔ19-LV particles and effector cells. 2.5×10^5^ effector cells transfected with 750 - 7.5 ng/T75 of S protein encoding plasmid were compared with three-fold dilutions of the particles starting with 2.5×10^9^ VLPs or 3 μL of LVs. Samples corresponding to the material used in the neutralization experiments are marked with arrowheads.

### Neutralization of the membrane fusion activities

Having established three different quantitative assays systems with an extraordinary high signal-to-noise ratio, we next assessed the neutralizing capacity of an S protein specific antibody, as well as convalescent sera from two donors. The sera were obtained four months after diagnosis of the infection with SARS-CoV-2 by PCR. Both donors had experienced mild symptoms. The strongest neutralization was observed in the particle entry assay. Addition of a neutralizing monoclonal anti-S antibody reduced the reporter signal by 2.3 orders of magnitude, which corresponded to more than 99% neutralization (Fig. 4A; Suppl. Fig. 3). Both sera reduced the reporter activity by 2.0 (98.5%) and 1.7 (97.4%) orders of magnitude, respectively, compared to control serum (Fig. 4A; Suppl. Fig. 3). In sharp contrast, their effect on cell-cell fusion remained below one order of magnitude for all three inhibitors (α-S AB: 0.4 orders (60.8%); serum 81: 0.5 orders (71.0%)); serum 82: 0.7 orders (79.1%)), even though the signal-to-background ratio was comparable for both assays (Fig. 4B; Suppl. Fig. 3). For the inhibition of FFWO, a mixed picture was obtained. The monoclonal anti-S antibody decreased FFWO signal by 1.7 orders of magnitude to background level (97.9% neutralization) (Fig. 4C). Titration of the antibody yielded an IC50 of 0.37 μg/mL, well in line with the value determined by the vendor in pseudovirus neutralization assays (0.11 μg/mL) (Fig. 4D). The effect of the patient sera was clearly less pronounced than what was observed in particle entry neutralization. The strongest effect was seen with serum 82 which decreased the fusion signal by 1.2 orders of magnitude (94% neutralization). Serum 81, which was highly active in the entry assay had no stronger effect on the fusion signal than the control serum (0.5 orders of magnitude, 72% neutralization) (Fig. 4C-D). Notably, antigen amounts on SΔ19-LV particles and the corresponding VLPs were similar, while a higher dose of S was used in the FFWO neutralization experiment than in the particle entry neutralization experiment (Fig. 4E).

They exceeded the amounts of S protein present in the cell-cell fusion assay by far, thus excluding higher amounts of antigen in the cell-cell fusion assay as causative for this difference.

## Discussion

Here we provide novel assay systems for a systematic side-by-side comparison of the membrane fusion activities of the SARS-CoV-2 spike protein S. Three quantitative assays are presented, covering i) entry of LV particles pseudotyped with S, ii) fusion between S and ACE2 expressing cells and iii) fusion between ACE2 expressing cells via VLPs displaying S protein. All assays exhibit high signal-to-noise ratios covering two to three orders of magnitude as important prerequisite to identify subtle differences, e.g. when screening for the activities of inhibitors. Several studies have recently published S protein based particle cell entry assays relying either on replication incompetent VSV particles (Hoffmann *et al.*, 2020b) or LV vectors similar to those used here in this study. With a maximum signal-to-noise ratio of four orders of magnitude the system we describe here is well among or even better than the top-performing LV-based assays published so far with second (Ou *et al.*, 2020) or simply first generation LVs with their significant safety concerns (Zeng *et al.*, 2020; Zhu *et al.*, 2020). It thus offers improved safety, as vectors are produced without the use of helper virus or separate packaging and transfer plasmids, significantly lowering the risk of replication-competent virus formation. Regarding cell fusion assays, previously published studies relied on nucleus counting or reporter complementation (Ou *et al.*, 2020; Zhu *et al.*, 2020). The assay system we describe here improves upon the previously reported maximum signal-to-noise ratios by at least an order of magnitude (Zhu *et al.*, 2020).

The syncytia forming activity of the S protein is remarkable not only with respect to speed and extent, but even more so with respect to the low amounts of S protein required, even when ACE2 is not overexpressed on the target cells. The high sensitivity of the assay we established here allowed the detection of cell fusion activity at levels of S protein which were below the level of detection of flow cytometry and Western blot. This remarkable activity is likely enabled by the high affinity of S protein for its receptor ACE2 (Shang *et al.*, 2020b)(Walls *et al.*, 2020). Being well in the low nanomolar range, it is even lower than that of the highly fusogenic MV for its receptors (Navaratnarajah *et al.*, 2008; Mühlebach *et al.*, 2011).

To our knowledge, we here provide the first demonstration of FFWO not only for the SARS-CoV-2 S protein but for any coronavirus, while FFWO has been observed for other enveloped viruses entering cells by pH-independent membrane fusion, such as retroviruses (Clavel and Charneau, 1994), paramyxoviruses (Bratt and Gallaher, 1969) and herpesviruses (Falke *et al.*, 1985). While it is conceivable that high concentrations of virus particles may accumulate in certain organs of infected patients, clear evidence for a pathogenic relevance of FFWO in patients has yet to be provided, although some evidence from animal studies exists for retroviruses (Murphy and Gaulton, 2007). Independently of that, the FFWO assay described here can become a prime choice for the preclinical evaluation of neutralizing binders for the following reasons: First, it can be performed at BSL-1 conditions, while LV-based assays require BSL-2. Second, it exhibits a high signal-to-noise ratio, even exceeding the maximum ratio recently reported for a benchmark LV-based neutralization assay (Zeng *et al.*, 2020). Third, this assay is unique in combining neutralization of particle-associated S protein with neutralization of cell fusion, thus providing information about two mechanisms of infection in a single readout.

Syncytia formation has recently been described as main and unique lung pathology in patients affected with Covid-19 in an occurrence not seen in other lung infections before. The polynucleated pneumocytes detected by histology and immunostaining were virus positive and expressed spike protein (Giacca *et al.*, 2020). Especially in persistent virus infections, syncytia formation is an important mechanism of virus spread as described for other important human pathogens like MV (Ferren *et al.*, 2019) or various herpesviruses (Cole and Grose, 2003). The low level of S protein sufficient for cell fusion we determined here appears to be ideal to support persistence of SARS-CoV-2. Evidence for SARS-CoV-2 persistence including neurological complications is recently accumulating and it is particularly the persistence which appears to result in fatal outcomes (Hamming *et al.*, 2004; Berger, 2020). In this respect, it may be worrying that the antibodies and convalescent sera we tested were less efficient in neutralizing cell-cell fusion than particle entry, even more so since the amounts of S protein present in the latter assay were substantially higher. This suggests that cell fusion is not only proceeding with minimal amounts of S protein, but also difficult to access for neutralizing antibodies. Further studies assessing sera from many more patients including severe and fatal cases will have to be performed to investigate this hypothesis further. The biological assay systems required are now available.

## Acknowledgement

The authors like to thank Klaus K. Conzelmann (Munich) for providing the plasmid encoding codon optimized spike protein. Additionally, we like to thank Gundula Braun and Manuela Gallet for excellent technical assistance in generating lentiviral vector stocks and cloning of plasmids

## Author contributions

S.T. designed, performed and evaluated experiments. A.M. designed experiments, analysed data and contributed to writing of the manuscript. V.R. designed and performed experiments and generated unique research material. T.J.M. provided unique research material. E.F., and K.C. contributed to conceiving the study and supervising work. C.J.B. conceived the study, supervised work, acquired grants and wrote the manuscript.

