## Supplemental Material for "Quantitative Assays Reveal Cell Fusion at Minimal Levels of SARS-CoV-2 Spike Protein and Fusion-from-Without"

**Short title: SARS-CoV-2 membrane fusion**

Samuel A. Theuerkauf[1*], Alexander Michels[1*], Vanessa Riechert[1], Thorsten J. Maier[2], Egbert Flory[3], Klaus Cichutek[1], and Christian J. Buchholz[1,3]

[1]Molecular Biotechnology and Gene Therapy, Paul-Ehrlich-Institut, Langen, Germany; [2]Division Safety of Medicinal Products and Medical Devices, Paul-Ehrlich-Institut, Langen, Germany; [3]Division of Medical Biotechnology, Paul-Ehrlich-Institut, Langen, Germany

[*]These authors contributed equally to this work.

Correspondence: Christian J. Buchholz, Molecular Biotechnology and Gene Therapy, Paul-Ehrlich-Institut, Paul-Ehrlich-Straße 51-59, 60528 Langen, Germany;, phone: +496103774011, fax: +496103771255.

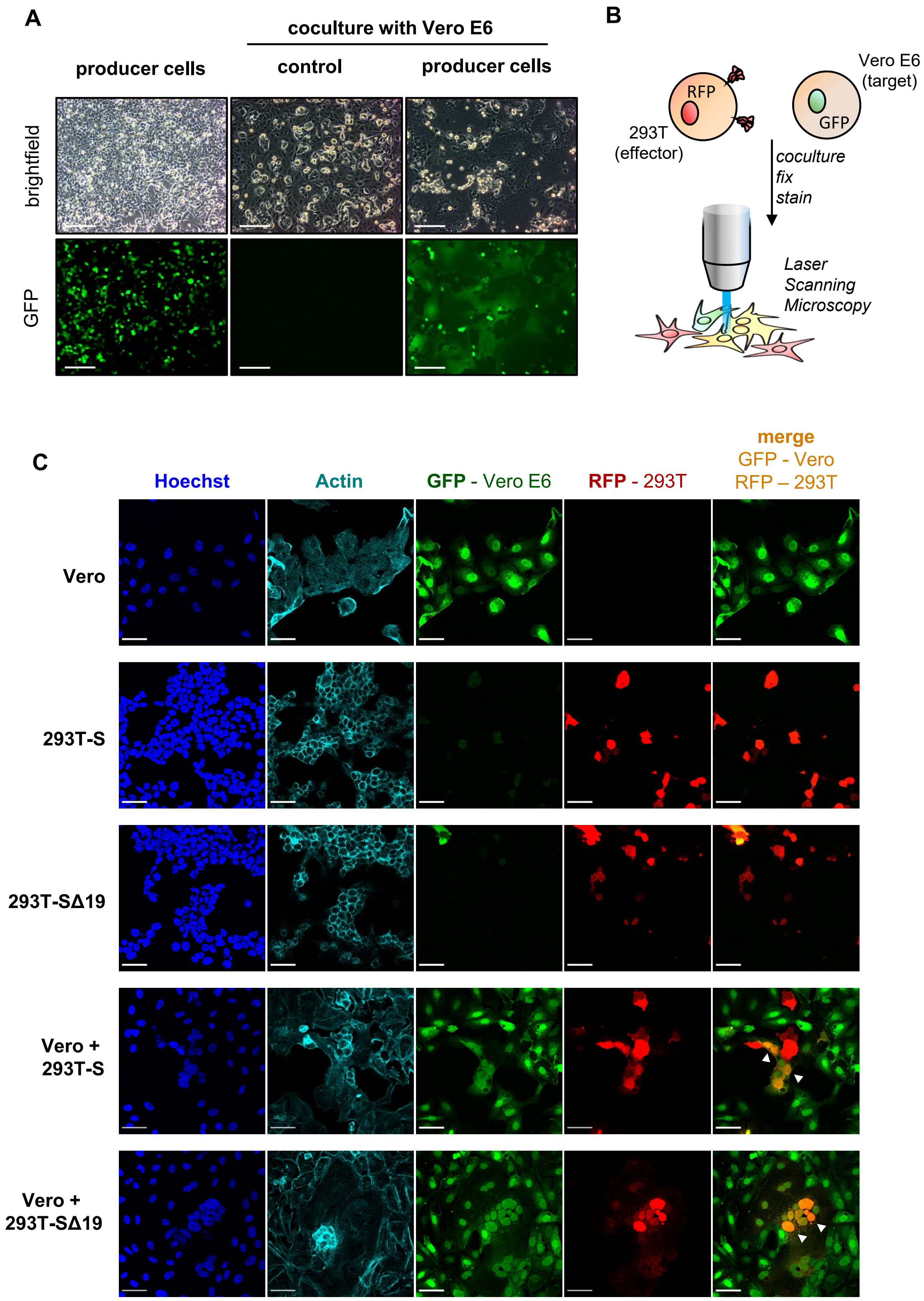
A
coculture with Vero E6
producer cells
control
producer cells
brightfield
GFP
B
RFP
293T
(effector)
Vero E6
(target)
GFP
coculture
fix
stain
Laser
Scanning
Microscopy
C
Hoechst
Actin
GFP - Vero E6
RFP - 293T
merge
GFP - Vero
RFP – 293T
Vero
293T-S
293T-SΔ19
Vero +
293T-S
Vero +
293T-SΔ19

**Figure S1: Microscopic evaluation of S protein mediated cell fusion.**

(**A**) HEK-293T packaging cells releasing SΔ19-LV having the GFP reporter gene packaged or untransfected control cells were detached by trypsinization and cocultured with Vero E6 cells. After overnight reattachment, presence of syncytia in the cocultures was determined by brightfield (top panel) and epifluorescense microscopy (bottom panel). Scale bars are 500 µm. (**B**) Workflow for the microscopical assessment of syncytia formation followed by color mixing induced by SARS-CoV-2 S. 293T cotransfected with RFP and SARS-CoV-2 S (FL or Δ19) were cocultured with Vero E6 cells stably expressing GFP. Cocultures were fixed and stained for nuclei (Hoechst) and actin (phalloidin) before being imaged by confocal laser scanning microscopy. (**C**) Confocal laser scanning micrographs of cocultures described in (B). Scale bars are 50 µm. Where necessary, micrographs underwent differential histogram stretching to ease qualitative analysis of cell morphology. Arrows point to signal colocalization resulting from fusion of both cell populations.

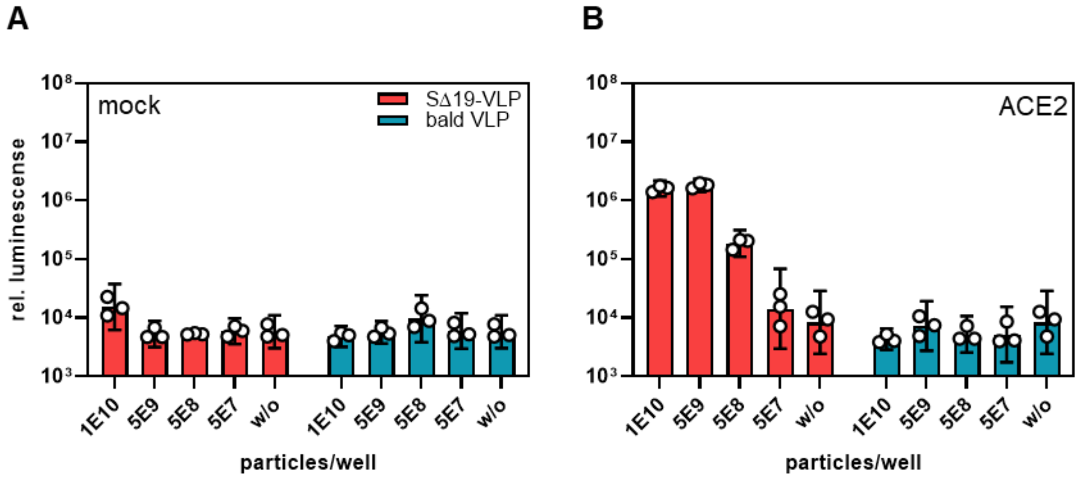


**Figure S2: Fusion-from-without is enhanced by ACE2**

The indicated numbers of SΔ19-VLP or bald VLP were added to cocultures of HEK-293T target cells expressing the α- and ω-fragments of β-galactosidase. In addition cells were transfected with a mock plasmid (**A**) or the ACE2 encoding plasmid (**B**). Reporter complementation was quantified in luminescence reactions after overnight incubation. Bars and error bars represent geometric means of technical triplicates and 95 % confidence intervals, respectively.

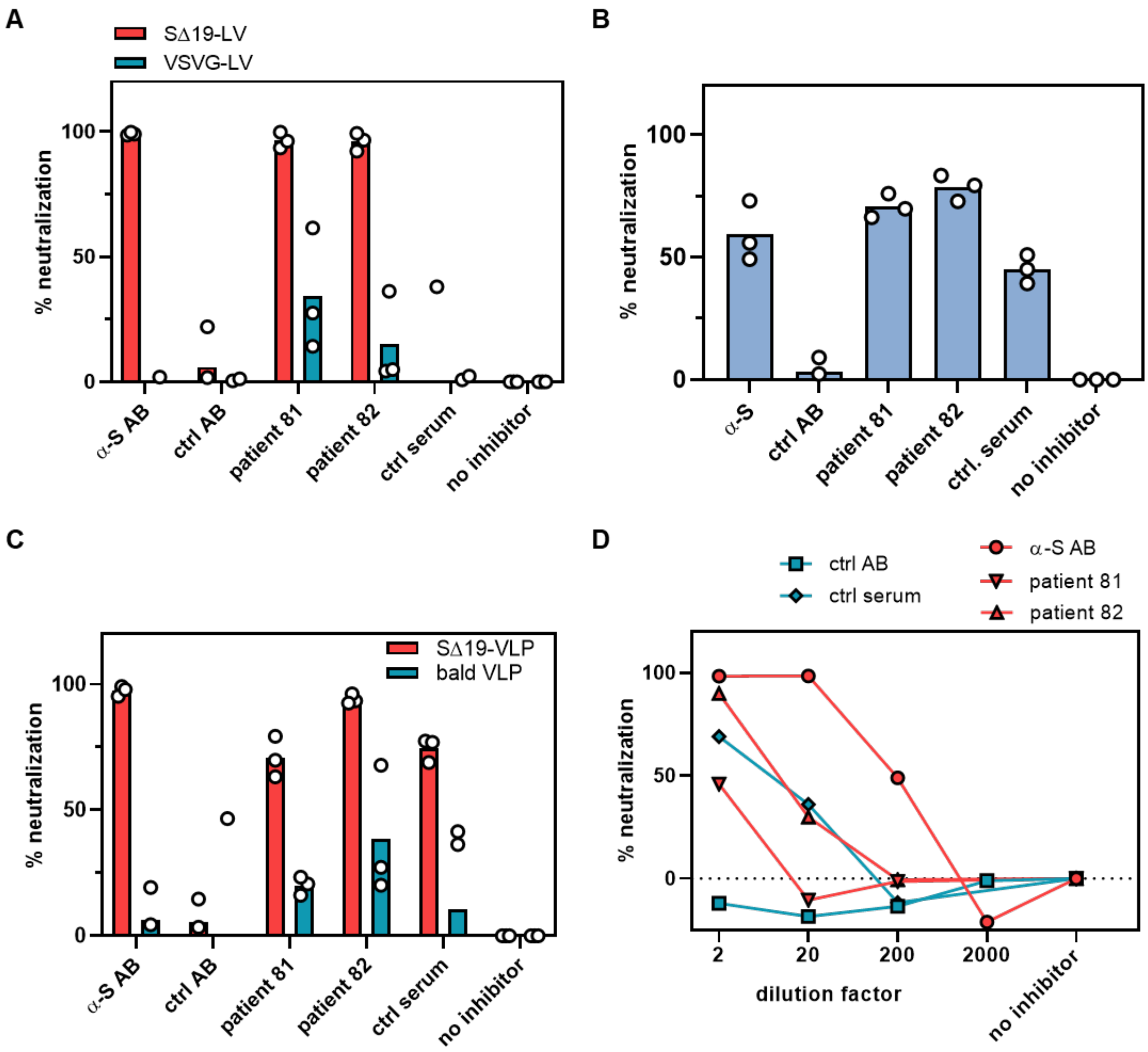


**Figure S3: Antibody mediated neutralization of membrane fusion**

Alternative representation of the data represented in Fig. 4A-D. The neutralizing activities of the S-protein specific antibody and the sera from two convalescent Covid-19 patients were determined against S-protein mediated particle entry (**A**), cell-cell fusion (**B**) and FFWO (**C-D**). Data is represented as percent neutralization relative to the control without any inhibitor, with each symbol representing the neutralization determined in a separate run. Bars represent means of neutralization activities.
